# Potential impacts of insect-induced harvests in the mixed forests of New England

**DOI:** 10.1101/692376

**Authors:** Meghan Graham MacLean, Jonathan Holt, Mark Borsuk, Marla Markowski-Lindsay, Brett J. Butler, David B. Kittredge, Matthew J. Duveneck, Danelle Laflower, David A. Orwig, David R. Foster, Jonathan R. Thompson

## Abstract

Forest insects and pathogens (FIPs) have significant impacts on U.S. forests, each year affecting an area nearly three times the area of wildfires and timber harvesting combined. We surveyed family forest owners (FFOs) in the northeastern U.S. and 84% of respondents indicated they would harvest in at least one of the presented FIP infestation scenarios. This harvest response represents a potentially significant shift in the timing, extent, and species selection of harvesting in the Northeast. Here we used the landowner survey, regional forest inventory data, and characteristics of the emerald ash borer (EAB, *Agrilus planipennis*) invasion to examine the potential for a rapidly spreading FIP to alter harvest regimes and affect regional forest conditions. Twenty-five percent of the FFO parcels in the Connecticut River Watershed in New England are likely to be harvested in response to EAB within 10 years. This prediction represents an increase in harvest frequencies, from 2.9%/yr (historically) to 3.7%/yr, on FFO woodlands. At typical harvest intensities, this would result in 13% of the total aboveground biomass removed through these harvests, with 81% of that biomass from species other than ash, creating a forest disturbance that is over twice the magnitude of the disturbance from EAB alone.

## INTRODUCTION

Forest insects and pathogens (FIPs) have significant impacts on U.S. forests, each year affecting an area nearly three times the area of all wildfires and timber harvesting combined (Hicke et al. 2012, Williams et al. 2016). FIPs selectively reduce tree species abundances, thereby altering forest structure and composition, disrupting carbon, water, and nutrient cycles (Hicke et al. 2012), and undermining ecosystem service provisioning, including timber production, carbon storage, and wildlife habitat creation (Boyd et al. 2013). FIPs also have significant indirect impacts on forests by altering management practices, often initiating pre-emptive and salvage harvesting (Lindenmayer et al. 2008). In 2017, our interdisciplinary team surveyed family forest owners (FFOs) in the Northeastern, U.S.; approximately 84% of respondents indicated that they would harvest their trees in at least one of the four presented FIP infestation scenarios (Markowski-Lindsay et al. 2019). This harvest response to FIPs represents a potentially significant shift in the timing, extent, and species selection of harvesting in the Northeast. Timber harvesting is the primary driver of forest disturbance in the northeast (Brown et al. 2018); therefore, this shift in the harvest regime would change regional patterns of disturbance with potentially profound impacts. Here we use the survey and regional forest inventory data to examine the potential for an emerging and rapidly spreading FIP, the emerald ash borer (*Agrilus planipennis*), to alter harvest regimes and affect regional forest conditions.

Salvage harvesting in response to natural disturbances, like FIPs, is commonly practiced to recover some of the value in their timber before it deteriorates (Lindenmayer et al. 2008). Like all harvests, salvage harvests can alter forest development trajectories and change structural legacies with lasting impacts on biodiversity and ecosystem services (Leverkus et al. 2018). Salvage harvests usually occur after the disturbance, but occasionally, and particularly in the case of FIPs, occur preemptively in an attempt to mitigate future damage or value loss, or slow FIP spread (Waring and O’Hara 2005). Here we use the term “FIP harvest” to refer to any harvest that was initiated either in anticipation of or in response to damage from a FIP. One concern with FIP harvesting in mixed hardwood forests, like those in the Northeast, is the phenomenon of harvest “by-catch”, or the non-host tree species harvested with the FIP-host species to enhance the commercial viability of the harvest or achieve another silvicultural objective (e.g. regeneration of a desirable mix of species). Irland et al. (1988) reported landowners harvesting to mitigate damages from spruce budworm (*Choristoneura fumiferana*) outbreaks in Maine, and those harvests included tree species beyond the budworm’s primary regional hosts. Similarly, Kizlinski et al. (2002) found that in harvests following hemlock woolly adelgid (*Adelges tsugae*) outbreaks in Connecticut and Massachusetts, species other than hemlock were removed in a majority of the harvests.

Northeastern forests are a carbon sink and are expected to remain so into the next century (Duveneck and Thompson 2019). However, the Northeast also has the largest abundance of exotic FIPs in North America and their numbers and impacts are predicted to increase with climate change and unchanged U.S. global trade policies (Lovett et al. 2016). The impacts of FIPs and FIP harvesting in the mixed forests of the Northeast threaten the status of these forests as a carbon sink and are increasingly important to understand, despite the inherent challenge of predicting landowner response to FIPs. In New England, FFOs control 54% percent of forested land (Butler et al. 2016), each with individual forest management and ownership objectives. Kittredge and Thompson (2016) found that the timing of harvests does not follow timber market trends, so most harvests on these woodlands are likely initiated by an exogenous force, such as a family event, change in income, or forest management plan. The results from our survey of landowners (Markowski-Lindsay et al. 2019), suggest that FIPs may serve as an important exogenous force for initiating harvesting in these woodlands, resulting in a regional disturbance regime that may be distinct from either the FIPs or typical harvesting alone.

The survey used a contingent behavior experiment to understand whether FFOs with parcels >4.05 ha (10 ac) in the Connecticut River Watershed (CTRW) of New England might harvest in response to a FIP (Markowski-Lindsay et al. 2019). Respondents were given four scenarios of FIP arrival and tree mortality and responded whether they would harvest in each scenario and provided a measure of certainty in their answer. The respondents of the survey fell into three categories: (1) Cutters (46% of respondents) – those that responded they would harvest for all four FIP scenarios; (2) Responsive Cutters (42%) – those that responded they would harvest in some of the scenarios presented; and (3) Non-cutters (12%) – those that would not harvest in any of the scenarios presented (Holt et al. 2019). Holt et al. (2019) then developed and defined agent functional types (AFTs) to characterize the different landowner responses to the presence of a generic FIP using the responses to the survey. Here we apply our knowledge of FFO response to FIPs and the developed AFTs to the specific case of emerald ash borer (EAB) to determine the potential direct and indirect impacts on the mixed forests of the CTRW.

EAB is a wood boring insect introduced from Asia that attacks North American ash (*Fraxinus* spp.). Since its identification in Michigan in 2002 (Herms and McCullough 2014), EAB has become the costliest exotic insect in the U.S. (Aukema et al. 2011). EAB typically kills ash trees in four to 10 years after infestation and has caused 99% mortality of ash trees > 2.5 cm in diameter in areas surrounding the initial introduction of EAB in Michigan (Klooster et al. 2018). EAB was first observed in New England in 2012 and the leading edge of the infestation is spreading at an average rate of 57 km/yr (Evans 2016), suggesting that EAB will be present throughout the region within the next decade (Herms and McCullough 2014).

Indeed, EAB has the potential to cause regional elimination of ecologically, economically, and aesthetically important hardwood species (Herms and McCullough 2014, Klooster et al. 2018). Information from loggers, foresters, and state agencies indicate that EAB is already prompting harvests and ash thinning (e.g., Mercader et al. 2015, Sadof et al. 2017). Ash often exists in relatively low abundance on forest woodlands (averaging 5.5% in New England, though patchy based on local conditions) (Bechtold and Patterson 2005). Therefore, given the volume of a typical harvest in the region and varied objectives of harvests, harvests initiated by the presence of EAB are likely to remove more than the affected ash. Here we seek to: (1) apply the results of a contingent behavior experiment to estimate a range of harvest responses to EAB based on the distribution of owner types and EAB invasion characteristics; and (2) quantify the resulting impact on regional forest biomass and composition.

## METHODS

### Study area and spatial data

We conducted our study in the Connecticut River Watershed (CTRW), a 3-million ha watershed within New Hampshire, Vermont, Massachusetts, and Connecticut (**Figure 1**). There are over 90,000 FFOs in the CTRW. We compiled ownership parcel boundaries for the CTRW from: state-level GIS sources, county and/or town websites, and tax maps. Where parcel maps were unavailable, we imputed parcel boundaries from neighboring towns with similar demographic and spatial distributions (approx. 12% of the landscape). A map of tree species aboveground biomass (AGB) was estimated following the methods in Duveneck et al. (2015) to associate each pixel with Forest Inventory Analysis (FIA, Bechtold and Patterson 2005) plots with similar spectral and environmental characteristics. We used this forest map to estimate the impacts of FIPs and subsequent harvest, as well as determine which tree species co-occur with ash and therefore may be most affected by the harvest response to EAB (Appendix A). For this study, we only used the distributions of white ash (*Fraxinus americana*) and black ash (*Fraxinus nigra*), and while these are the two most abundant ash species in the CTRW, white ash makes up 99.96% of the total ash biomass (Table 1).

**Table 1.** Merchantable timber aboveground biomass (AGB) of the 10 most abundant species and two most abundant ash species* in harvestable FFO parcels in the CTRW and the % harvested in response to EAB for each scenario.

| Species | Total AGB<br>(Tg) | % of total AGB harvested |  |  |
| --- | --- | --- | --- | --- |
|  |  | Scenario A: typical<br>intensity, equal<br>removal | Scenario B: typical<br>intensity and<br>preference | Scenario C: decreased<br>intensity, increased<br>preference |
| *White ash (*Fraxinus americana*) | 8.2 | 54%±2.4% | 54%±2.4% | 54%±2.4% |
| *Black ash (*Fraxinus nigra*) | 3.6x10<sup>-3</sup> | 52%±5.7% | 52%±5.7% | 52%±5.7% |
| Red maple (*Acer rubrum*) | 27.1 | 11%±0.6% | 11%±0.6% | 5%±0.3% |
| White pine (*Pinus strobus*) | 26.4 | 11%±0.7% | 9%±0.6% | 4%±0.3% |
| Sugar maple (*Acer saccharum*) | 26.4 | 13%±0.6% | 11%±0.5% | 5%±0.2% |
| Red oak (*Quercus rubra*) | 22.8 | 10%±0.8% | 9%±0.7% | 4%±0.3% |
| Eastern hemlock (*Tsuga canadensis*) | 19.8 | 11%±0.7% | 9%±0.6% | 4%±0.3% |
| Yellow birch (*Betula alleghaniensis*) | 8.4 | 13%±0.8% | 17%±1.0% | 9%±0.5% |
| American beech (*Fagus grandifolia*) | 7.0 | 12%±0.6% | 13%±0.7% | 6%±0.3% |
| Paper birch (*Betula papyrifera*) | 4.6 | 12%±0.7% | 16%±1.0% | 9%±0.6% |
| Black cherry (*Prunus serotina*) | 4.2 | 11%±0.8% | 14%±1.0% | 8%±0.6% |
| Black birch (*Betula lenta*) | 4.1 | 9%±0.8% | 11%±1.0% | 6%±0.6% |
| **Total** | **180.2** | **13%±0.8%** | **13%±0.8%** | **7%±0.5%** |

**Figure 1.**
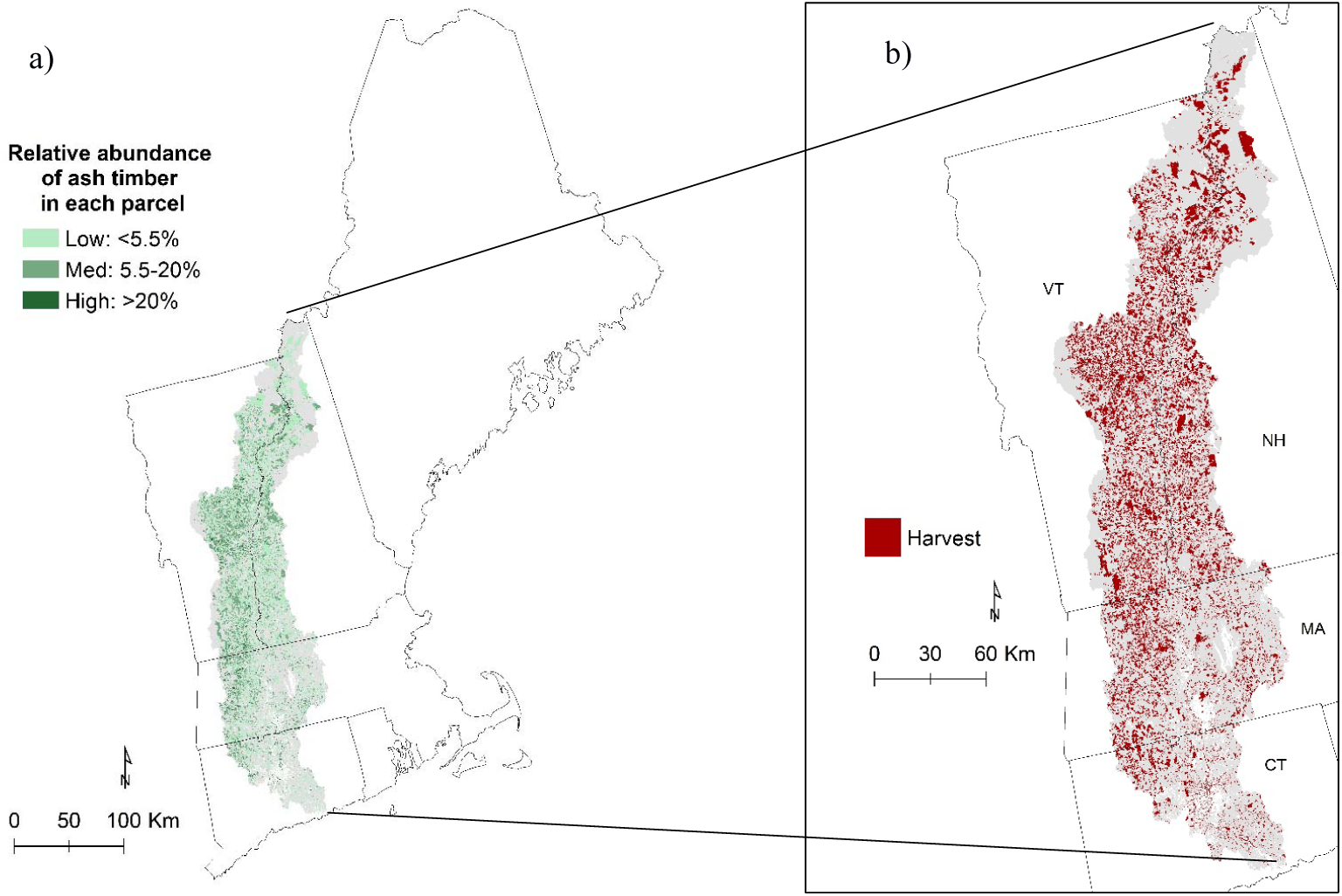
a) Relative abundance of merchantable ash timber volumes for each pixel in the CTRW of New England. Grey areas represent non-FFO parcels or parcels without ash. b) An example harvest map where grey areas are parcels not selected for harvest in this replicate.

### Determining harvest response to EAB

In this region, a majority of FFO harvests are low intensity (e.g. non-commercial) harvests, but are ecologically important drivers of forest change (Thompson et al. 2017, Belair and Ducey 2018). Therefore, for modeling the harvest response to EAB, we assumed the range of possible harvest responses included commercial and non-commercial harvests. We also assumed that EAB was present throughout the study area (at its current rate of spread, this assumption will likely be met within one decade). Following Holt et al. (2019), each parcel with ash was assigned an AFT and a probability of harvest based on the presence of EAB (Figure 2, and Appendix B for details). For the Cutter and Non-cutter AFTs, harvest probabilities were assigned using the certainty distributions of the survey responses from the respondents in these AFTs (e.g., 1 = Cutter with high certainty; 0 = Non-cutter with high certainty), where the certainty distributions were derived from a 5-step Likert scale certainty question that followed each scenario in the survey (Appendix B). The probability of harvest for Responsive Cutters was based on what Holt et al. (2019) found to be significant in predicting harvest for this group: 1) the percentage of their trees killed by the FIP; and 2) the time it takes for those trees to die (Figure 2). A higher mortality percentage and more rapid tree death both increased the likelihood a Responsive Cutter would harvest. For this EAB application, the relative abundance of ash on the parcel was used as the mortality percent and we made the conservative assumption of 8 years to mortality from initial infestation for those ash trees. The probabilities of harvest were then stochastically used to predict which parcel owners might harvest in response to EAB. The process of assigning AFTs, probabilities of harvest, and finally harvest locations, was replicated 10 times to capture the variability in assigning harvest locations.

**Figure 2.**
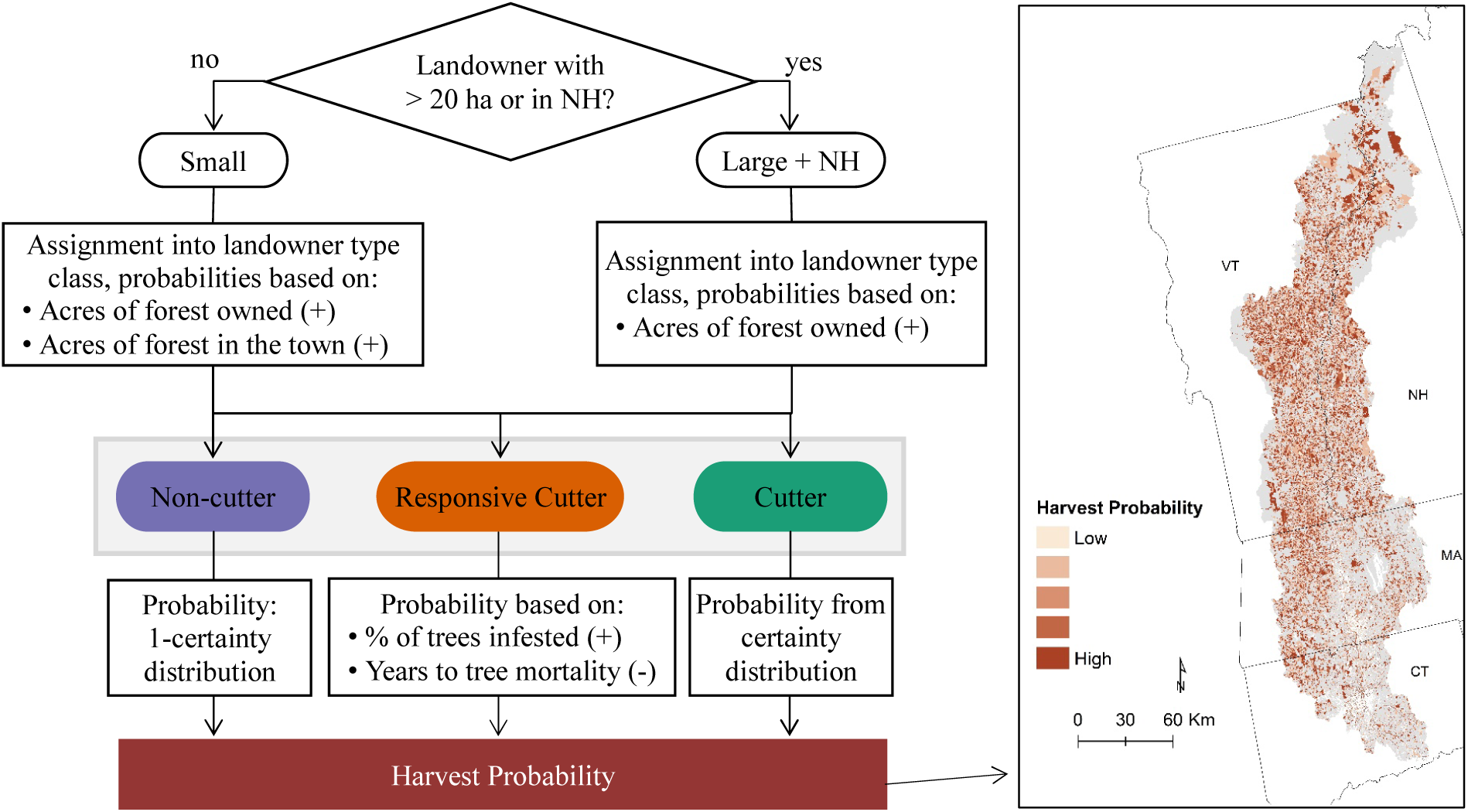
We followed the methods from Holt et al. (2019) to assign AFTs to the FFO parcels in the CTRW. We then used the information about the AFTs, survey response certainty distributions, and specifics from the EAB invasion to calculate harvest probabilities for each of the parcels. The relationships between probability of being classified as a Cutter landowner type or with higher probability of harvest are shown in parenthesis. Grey areas represent non-FFO parcels or parcels without ash.

### Quantifying cumulative impacts of EAB

We used the forest composition map and the 10 simulated harvest maps to estimate species AGB removal in EAB-initiated FFO harvests. To account for the uncertainty regarding the specific silvicultural choices made by foresters and loggers, we examined three alternative harvest scenarios and report the total AGB and species-specific AGB that could be harvested in response to EAB. Our estimates include the AGB of merchantable timber trees (i.e., larger than 23 cm (9 in) diameter at breast height (dbh) for softwoods and 28 cm (11 in) dbh for hardwoods) and only parcels that met the criteria for our survey (i.e., FFO parcels > 4.05 ha). We also set a minimum forest area size of 3.15 ha on mixed land use parcels to ensure enough forest for measurable tree removal. We included three harvest scenarios, described below.

#### Scenario A: Typical harvest intensities and no species preferences

Scenario A assumed that the presence of a FIP is an exogenous force that initiates a typical FFO harvest, rather than a reactionary salvage harvest. As some state agencies and interest groups recommend removing healthy ash just prior to infestation (e.g., see NHBugs.org), some landowners are choosing to pre-emptively harvest ash, which may indeed result in a more planned and typical harvest than a reactionary harvest after infestation. Furthermore, the harvests in this scenario did not differentially affect tree species based on their type or value; rather, the proportion of AGB removed, or harvest intensity, was applied equally to all species with trees of merchantable timber size on the parcel. To determine what constitutes a “typical harvest” intensity for FFO harvests in the CTRW, we used the harvest intensity distributions from re-measured FIA plots on FFO lands in the CTRW states that experienced tree removal between measurements. We included all types of removals, from very low intensity removals to clear-cuts, to define these distributions (Appendix B). Harvest intensities for individual parcels were then drawn from these distributions. All merchantable ash was first removed, then any remaining timber was removed proportionally from all remaining species until the target harvest intensity was met. Where the total volume of ash on a parcel exceeded the necessary volume for the target harvest intensity, all merchantable ash was removed, but no more.

#### Scenario B: Typical harvest intensities and species preferences

Scenario B differentially harvests tree species based on patterns observed in the historical data. In this scenario, the harvest intensities were the same as in Scenario A, but the amount removed for each species (other than ash) was weighted to match the observed harvest distributions from the re-measured FIA plots. Weights were calculated by dividing the proportion of each species removed from the initial measurement to the re-measurement of a plot by the overall harvest intensity for that plot and then averaged over all harvested plots (Appendix C).

#### Scenario C: Decreased harvest intensity and increased species preference

In Scenario C, we examined the possibility that harvest prompted by EAB will be less intense than typical harvests, supposing that many of the harvests were reactionary and not planned with harvest volumes as an objective. Therefore, the harvest intensities for each parcel were reduced by 50%. In addition, the weighting of species removal was intensified to approximate a behavior where preferable trees are selectively chosen in a low-intensity FIP harvest so the landowner can recover the operational cost of harvesting while removing the fewest number of trees beyond the affected ash. To reweight the species, weights of preferential trees (weight > 1) were increased by 10% and the weights of non-preferential trees (weight < 1) were decreased by 10%.

## RESULTS

### Determining harvest response to EAB

In total, 80,010 FFO parcels met the area requirements for harvest consideration, and of these parcels, 40,419 contained merchantable ash timber. On average, 42%±0.2% (mean±sd) of the parcels with ash were assigned to the Cutter landowner type class; 44%±0.1% were Responsive Cutters, and; 14%±0.1% were Non-cutters. The average probability of harvest for Cutters was 0.89±0.003, 0.22±0.004 for Responsive Cutters, and 0.12±0.004 for Non-cutters (Figure 2). On average, 25%±0.07% (19,705±55 parcels) of the FFO parcels that met the area requirements for harvesting (with or without ash) were harvested, equating to 37%±0.7% of the forested area of all FFO parcels in the watershed. Approximately 77%±0.3% of the harvests occur on Cutter type class parcels, 19%±0.3% on Responsive Cutter parcels, and 4%±0.1% on Non-cutter parcels (Figure 1). Harvest intensities also vary by location within the CTRW (Appendix B). The average harvest intensity applied to harvest parcels in the northern region was 45%±28%, and the average harvest intensity applied to the southern region was 27%±22%.

### Quantifying cumulative impacts of EAB

We found 8.2 Tg of merchantable timber size ash biomass on harvestable FFO parcels in the CTRW; up to 99% of which is expected to be killed by EAB (Klooster et al. 2018).

Approximately 54% (4.4 Tg) of that ash is expected to be removed from the landscape through harvest in all scenarios (Table 1). In Scenario A, 18.8 Tg of AGB was harvested from co-occurring species (or as by-catch) in addition to the ash (Table 1). Including the ash removal, the total AGB harvested in response to EAB equates to 13% of all AGB in the harvestable FFO parcels in the CTRW (with or without ash). In Scenario B, a similar amount of biomass was removed as in Scenario A; however, the amount of each species removed differed. In this scenario, yellow birch (*Betula alleghaniensis*), paper birch (*B. papyrifera*), and black cherry (*Prunus serotina*) incur the largest potential losses, up to 17% of their AGB, due to preferential harvesting and co-occurrence with ash (Table 1 and Appendix A). In Scenario C, with decreased harvest intensity only 9.0 Tg of biomass was harvested from co-occurring species. However, a few species, such as yellow birch, were removed at higher than average proportions.

## DISCUSSION

We used forest owner survey data and a landowner typology to examine three scenarios describing the harvest response to EAB; they suggest that harvest frequency could increase 28% above the recent trends in harvesting (sensu Thompson et al. 2017). Twenty-five percent of parcels, representing 37% of the FFO forested area, within the CTRW were predicted to be harvested in response to EAB. Given the spread rate of EAB and its current locations within the watershed, most of this harvest is predicted to take place in the next decade. Assuming this future, the annual probability of harvest in response to EAB could be 3.7% for FFO lands in the CTRW for the next 10 years. Thompson et al. (2017) found that in the 20-state northeastern region of the FIA, the annual probability of harvest on non-corporate private woodlands was 2.9%. Therefore, the harvest induced by EAB likely represents an acceleration and potential expansion of harvest into parcels not usually harvested in this region.

At typical harvest intensities (e.g., scenarios A and B), 13% of the total AGB in forests in the CTRW could be harvested by FFOs in response to the invasion of EAB. In scenarios A and B, 81% of the removed AGB was from species other than ash (*Fraxinus* spp.). Therefore, the FIP harvest response in these scenarios created a disturbance, in terms of total AGB affected, more than twice the magnitude of the expected disturbance of EAB and included a larger number of species. In scenario C, where harvest intensities are lower, 65% of the removed biomass was from species other than ash, creating a disturbance 1.5 times the magnitude of EAB alone. Both co-occurrence with ash and silvicultural prescription influenced the scope of the disturbance for individual species. For example, 81% of yellow birch (*B. alleghaniensis*) is co-located with ash. Since yellow birch is also a preferred species for harvest, 17% of the merchantable timber-size yellow birch trees were harvested in the typical harvest intensities and species preference scenario (B). If average harvest intensities are lower for the FIP harvests but target preferential species (e.g., Scenario C), the cumulative impacts of EAB is less but preferred merchantable species that co-occur with ash (e.g. yellow birch) are removed proportionally more than species of less value or that do not co-occur with ash as frequently (e.g. black birch, *B. lenta*).

In all scenarios, more frequent harvests have far-reaching ecological impacts. Harvesting in response to a FIP with a specific host(s) alters the species composition of the landscape by differentially impacting species that co-occur with the host. The by-catch in FIP harvesting will be especially prevalent in the mixed forests of the Northeast when harvests are planned with timber volumes and/or larger regeneration openings as objectives, since, one host species often will not be enough to meet target volumes or silvicultural objectives.

### Limitations and future research

The survey we used to develop and apply the landowner typology did not ask questions about the type or intensity of harvesting that a FIP might induce. To account for this uncertainty, we presented three scenarios of varying harvest intensity and species removal. Further investigation into the intensity and types of silvicultural prescriptions applied in the harvest response to EAB is necessary to quantify these effects, particularly from the viewpoint of the foresters and loggers who are completing the harvests. Additionally, it is unknown if the harvest response may be limited by the capacity of foresters, loggers, and markets. We also expect that there is some interplay between harvest unrelated to EAB, FIP harvest in response to EAB, forest growth dynamics, and climate change. To assess specifically how FIP harvests may accelerate and intensify forest disturbance and change forest growth and composition over time, future research should explore these dynamics with a spatially and temporally explicit representation of background harvest and FIP harvest initiated by EAB.

In addition, there are several other FIPs currently impacting forests in the region (e.g., hemlock woolly adelgid, European gypsy moth (*Lymantria dispar*), beech bark disease), and Leung et al. (2014) predict a tripling of non-native wood-borers in the northeast by 2050, with at least one new wood-borer having the same or larger impact on the landscape as EAB. The interactions among FIPs will have extensive ecological impacts on the forested landscape, but as shown in this study, the coupled human management response may intensify and expand the total disturbance created by these FIPs. The individual ecological impacts of these FIPs on current forest structure and future forest dynamics is somewhat understood, however the management responses to multiple interacting global change drivers is less well-resolved. Our study underscores the importance of better understanding these complex interactions to predict the fate of our forests and their ability to continue to provide essential ecosystem services.

## ACKNOWLEDGEMENTS

Thank you to both Paul Catanzaro and Tony D’Amato for your thoughtful reviews of an early draft of this manuscript. This research is funded by the National Science Foundation Coupled Natural and Human Systems Grant No. DEB-1617075. This research is also partially supported by the Harvard Forest Long Term Ecological Research Program Grant No. NSF-DEB 12-37491.

## APPENDICES

### Appendix A: Co-occurrence of species with ash

The forest composition map was used to assess which species co-occur with ash on all FFO parcels.

**Table A1.** Tree species that most commonly co-occur on parcels with ash (*Fraxinus* spp.). Total merchantable timber is the total amount of species-specific aboveground biomass (AGB) in the CTRW. Average co-occurrence is 74%. The percent of total AGB that co-occurs with ash represents the amount of that species that occurs on parcels with ash; e.g., 84% of all black cherry in the CTRW occurs on parcels with ash.

| Species | Total merchantable timber AGB (Tg) | % of total AGB co-occurring with ash |
| --- | --- | --- |
| Black cherry (*Prunus serotina*) | 4.4 | 84% |
| American basswood (*Tilia americana*) | $1.7 \times 10^{-1}$ | 83% |
| Sugar maple (*Acer saccharum*) | 26.8 | 83% |
| Balsam poplar (*Populus balsamifera*) | $1.2 \times 10^{-2}$ | 82% |
| American beech (*Fagus grandifolia*) | 7.2 | 82% |
| Yellow birch (*Betula alleghaniensis*) | 8.4 | 81% |
| Paper birch (*Betula papyrifera*) | 4.6 | 80% |
| Bigtooth aspen (*Populus grandidentata*) | $8.0 \times 10^{-1}$ | 80% |
| Gray birch (*Betula populifolia*) | $2.6 \times 10^{-3}$ | 78% |
| Quaking aspen (*Populus tremuloides*) | 1.7 | 78% |
| Red maple (*Acer rubrum*) | 28.0 | 77% |
| Black birch (*Betula lenta*) | 4.1 | 76% |
| Eastern hemlock (*Tsuga canadensis*) | 20.2 | 76% |
| American hophornbeam (*Ostrya virginiana*) | $6.8 \times 10^{-2}$ | 76% |
| Red spruce (*Picea rubens*) | 3.6 | 75% |

### Appendix B: Process for assigning a harvest to each parcel

Following the methods from Holt et al. (2019), landowners were first categorized into two groups based on size and location of the parcel into a large parcel size + New Hampshire group and a small parcel size group (Figure 2). Then using a multinomial regression, landowners were assigned to one of the three AFTs: Non-cutter, Responsive Cutter, or Cutter. Moving beyond the AFT assignment from Holt et al. (2019), parcels were then given a probability of harvest unique to this example using EAB. Parcels in the Cutter or the Non-cutter AFTs were initially given 1 or 0 probability of harvest (respectively) and then those probabilities were augmented using the distribution of certainty of harvest from the survey responses of those groups (Figure B1). For example, a Cutter respondent that consistently ranked their responses as “Very Certain” would have a certainty of harvest of 1, while another Cutter respondent with lower certainty in their responses would have a certainty of harvest closer to 0.5. Likewise, a Non-cutter respondent that consistently ranked their responses as “Very Certain” would have a certainty of harvest of 0, while another Non-cutter respondent with lower certainty in their responses would have a certainty of harvest closer to 0.5. Cutter parcels were assigned individual harvest probabilities using a stochastic application of the Cutter certainty distribution and Non-cutter parcels were assigned individual harvest probabilities using 1-the Non-cutter certainty distribution. Holt et al. (2019) found that the probability of harvest for Responsive Cutters was based on the percentage of trees dying from the FIP on their land and the years to mortality for those trees, in which a higher mortality percentage and quicker time to death both increased the likelihood of harvest (Figure 2). For this specific application using EAB, the mortality percent was the relative abundance of ash for each parcel and we chose a conservative time to death of 8 years for ash at the time of initial infestation. The distribution of harvest probabilities was unique for each landowner type class (Figure B2). The probability of harvest was then used to stochastically assign harvest or no harvest to each parcel (Figure 1).

Subsequently, to mimic typical harvest intensities across the landscape, we assigned a harvest intensity (AGB removed/AGB prior to harvest) to each harvest parcel. We used remeasured FIA plots from the CTRW states to determine typical levels of harvest. An exploration of the harvest intensities in different regions and states where the CTRW resides revealed that the harvest intensities were different between Northern Vermont and Northern New Hampshire (as defined in the FIA state data) and the rest of the CTRW. The average harvest intensity of the northern region (N) was 46% of total AGB removed per harvest (n = 91), and the southern region (S) had an average harvest intensity of 27% (n = 120). The distributions of the harvest intensities captured by the FIA were also different. Therefore, multiple distributions to model the harvest intensity were tested (e.g. gamma, Weibull, lognormal, Cauchy, exponential), and the best fit was determined using the Akaike information criterion (AIC; Akaike 1973) for each region independently. For the northern region, a Weibull distribution (shape = 1.49, scale = 0.512) was the best fit for harvest intensities observed in the FIA (AIC = 26.82, ΔAIC = 2.16). For the southern region, a lognormal distribution (mean = −1.61, sd = 0.831) was the best fit for the distribution of harvests in that region (AIC = −86.70, ΔAIC = 1.66) (Figure B3). The process from AFT assignment to harvest map with harvest intensities was completed 10 times to create replicates for assessing the variability introduced in the process of determining harvest parcels and intensities.

**Figure B1.**
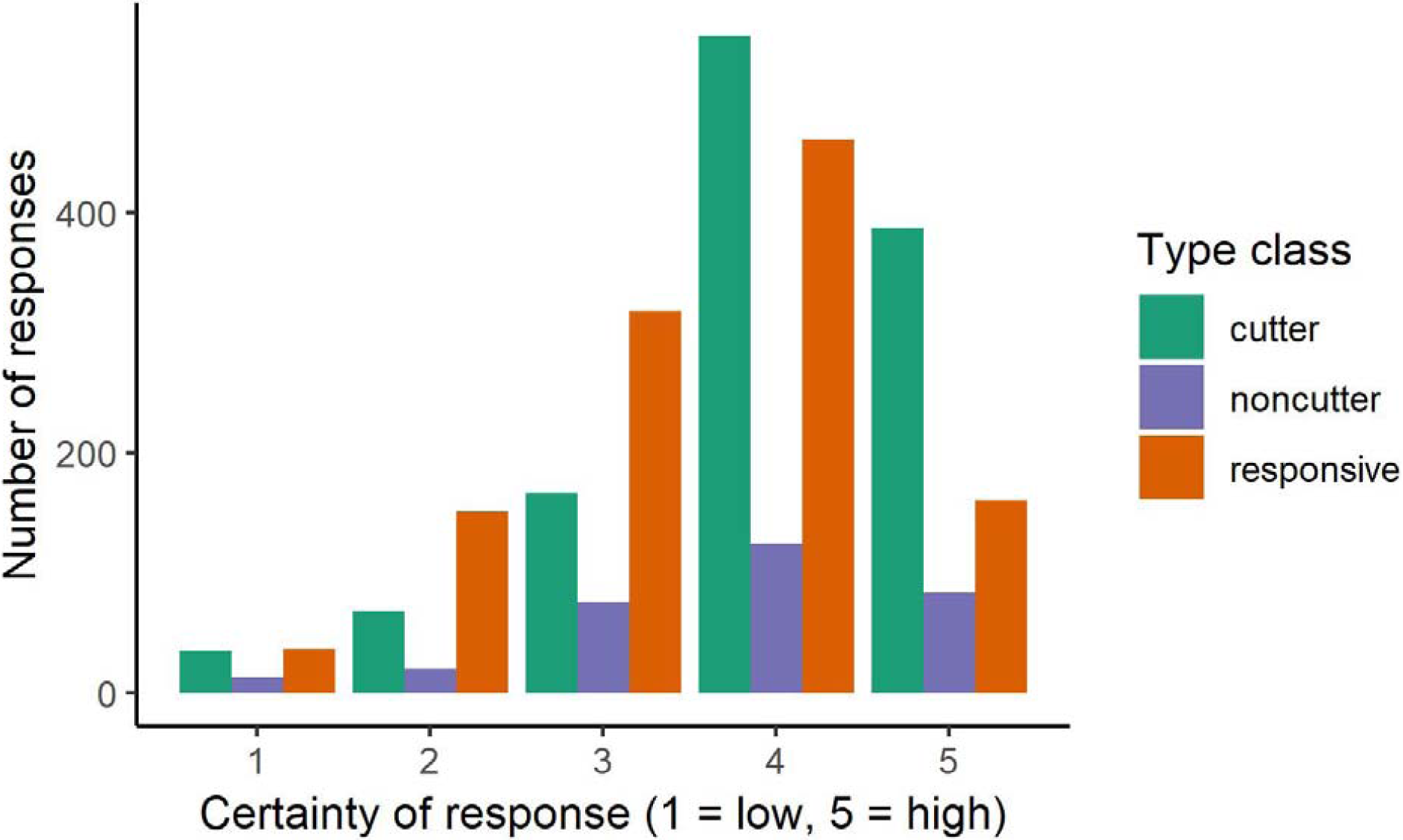
Certainty of responses by respondent category. The number of responses is the total number of responses to a scenario rather than responses by an individual landowner since certainties were unique to each scenario presented.

**Figure B2.**
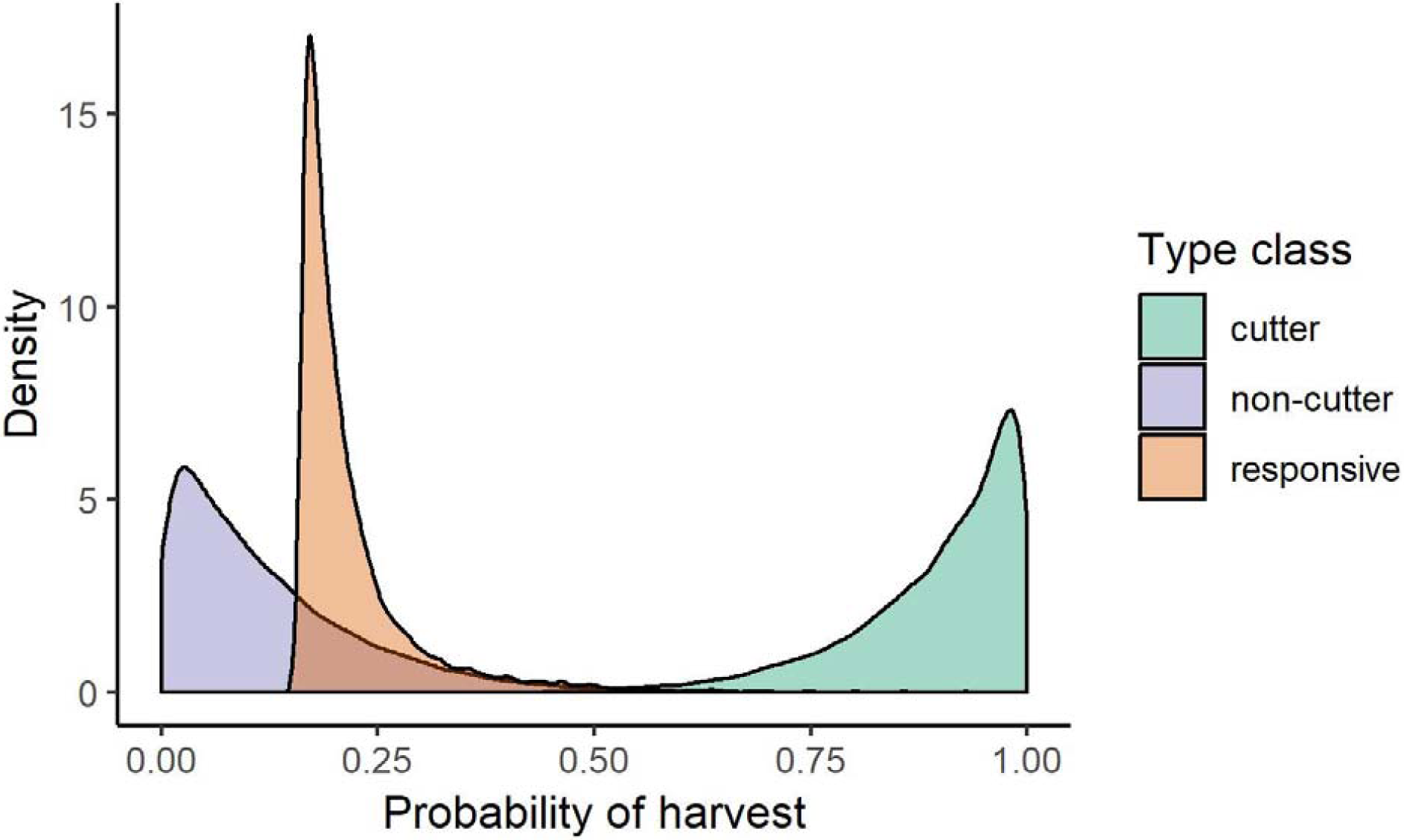
An example of the probability of harvest distributions for each of the land owner type classes for one of the 10 replicates. All replicates showed similar distributions.

**Figure B3.**
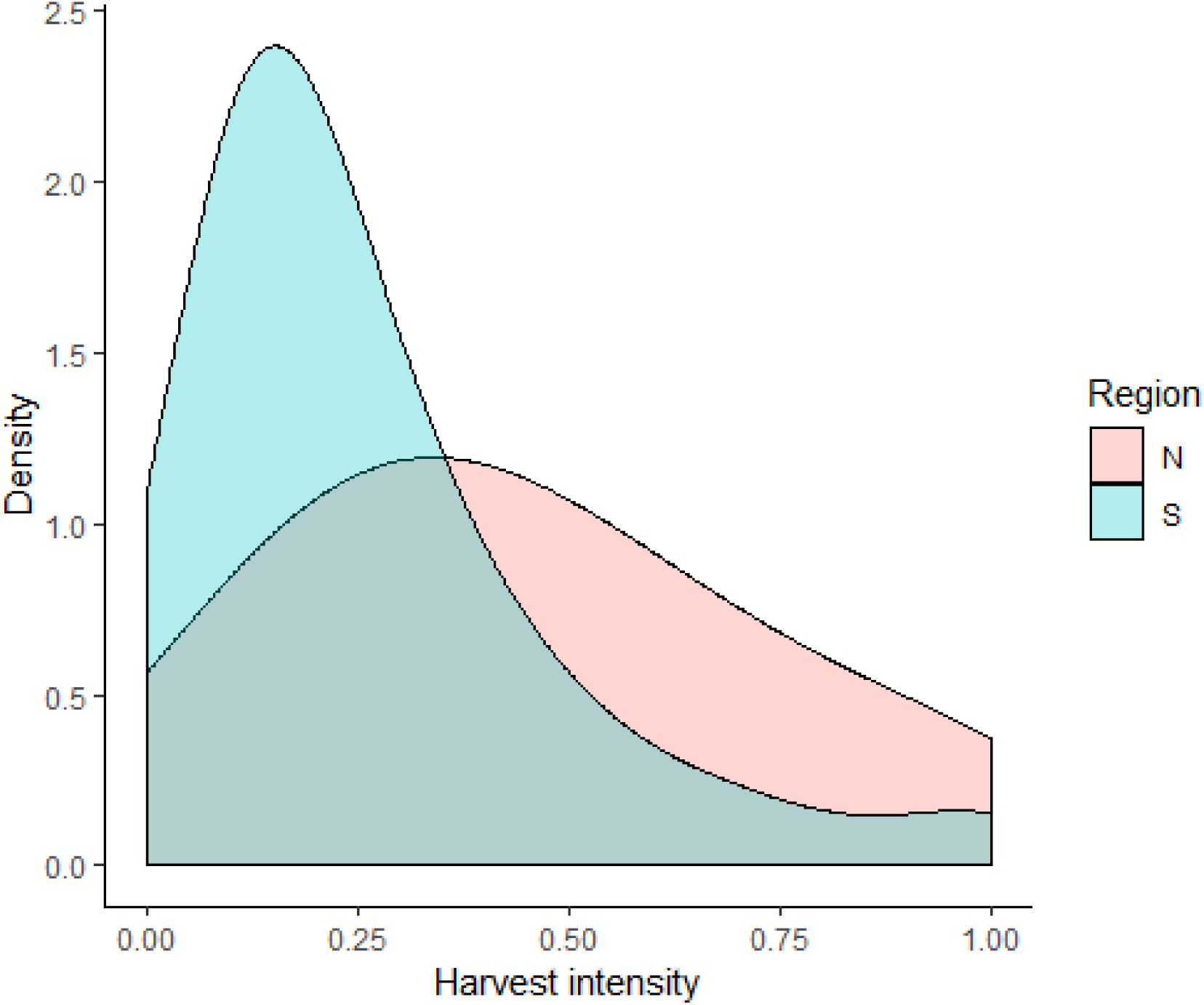
An example of the distribution of harvest intensities for each of the regions (N – Northern Vermont and Northern New Hampshire from the FIA state data, S – all other area in the CTRW states) for one of the 10 replicates. All replicates showed similar distributions.

### Appendix C: Weighting of species based on FIA species weights

Weights were based on species removals from individual FIA plots in the states of the CTRW. Weights were calculated by dividing the proportion of each species removed from the initial measurement to the re-measurement of each FIA plot by the harvest intensity of that plot. These proportions were then averaged over all harvested plots. Species removed in higher proportions than the average received a weight > 1, while species removed in lower proportions received a weight < 1. For example, if sugar maple (*Acer saccharum*) existed on a harvested plot, less of it was harvested than yellow birch (*Betula alleghaniensis*).

**Table C1.** The top ten most abundant species and their harvest weights as computed using remeasured FIA plots in Connecticut, Massachusetts, New Hampshire, and Vermont.

| Species | Weight |
| --- | --- |
| Red maple (*Acer rubrum*) | 0.95 |
| White pine (*Pinus strobus*) | 0.81 |
| Sugar maple (*Acer saccharum*) | 0.77 |
| Red oak (*Quercus rubra*) | 0.87 |
| Eastern hemlock (*Tsuga canadensis*) | 0.73 |
| Yellow birch (*Betula alleghaniensis*) | 1.24 |
| American beech (*Fagus grandifolia*) | 0.97 |
| Paper birch (*Betula papyrifera*) | 1.28 |
| Black cherry (*Prunus serotina*) | 1.08 |
| Black birch (*Betula lenta*) | 1.05 |

